# Robust, real-time and autonomous monitoring of ecosystems with an open, low-cost, networked device

**DOI:** 10.1101/236075

**Authors:** Sarab S. Sethi, Robert M. Ewers, Nick S. Jones, C. David L. Orme, Lorenzo Picinali

## Abstract

1. Automated methods of monitoring ecosystems provide a cost-effective way to track changes in natural system’s dynamics across temporal and spatial scales. However, methods of recording and storing data captured from the field still require significant manual effort.
2. Here we introduce an open source, inexpensive, fully autonomous ecosystem monitoring unit for capturing and remotely transmitting continuous data streams from field sites over long time-periods. We provide a modular software framework for deploying various sensors, together with implementations to demonstrate proof of concept for continuous audio monitoring and time-lapse photography.
3. We show how our system can outperform comparable technologies for fractions of the cost, provided a local mobile network link is available. The system is robust to unreliable network signals and has been shown to function in extreme environmental conditions, such as in the tropical rainforests of Sabah, Borneo.
4. We provide full details on how to assemble the hardware, and the open-source software. Paired with appropriate automated analysis techniques, this system could provide spatially dense, near real-time, continuous insights into ecosystem and biodiversity dynamics at a low cost.

## 1 Introduction

Scientists are increasingly moving towards automated methods of ecosystem monitoring: the need for such monitoring is pressing (Pereira and Cooper, 2006) and automatic data collection both helps avoid temporal and spatial undersampling and mitigates observer biases (Foster and Harmsen, 2012; Leach et al., 2016; Zwart et al., 2014). Advances in ecoinformatics and the processing of big data, such as through machine learning methods, means that we are well placed to analyse bulk data, but progress on automated recording to provide that data has been relatively restricted.

Two key barriers are providing a long-term power source and enabling automated data transmission. The field of acoustic monitoring (Acevedo and Villanueva-Rivera, 2006; Pijanowski et al., 2011) provides an example of early adoption of automatic methods and approaches to overcoming those barriers. One solution is to deploy commercial or open-source battery powered recorders (Hill et al., 2018; Maina et al., 2016; Whytock and Christie, 2016). These typically offer a few hundred hours of continuous recording but require regular visits post-deployment to collect data and replace batteries, increasing their effective cost and limiting their potential scalability. In contrast, more truly-autonomous systems employ solar panels to provide power and remotely upload the audio data, but have high unit costs and, in addition, need expensive communication infrastructure. For example, Cyberforest (Saito et al., 2015) uses expensive satellite data upload, and ARBIMON (Aide et al., 2013) requires a specialist radio data network.

Here we present an inexpensive autonomous ecosystem monitoring device based around a Raspberry Pi and the use of mobile data networks to implement continuous data collection from the field. Sensor configurations use a very general framework – we provide examples for a generic device, a time-lapse camera and continuous audio recording – facilitating long-term continuous monitoring from a variety of sensors with only minor modifications.

To realise truly-autonomous monitoring our method relies upon an existing mobile network in the study area, and access to direct sunlight for the solar power system. Despite these limitations, the system is designed with redundancy and is able to operate seamlessly over temporary inclement weather and disruptions to the mobile network. We have demonstrated its utility in real-world ecological surveys by deploying a network of monitoring units in the challenging environment of the tropical forests of Borneo.

## 2 Description

The unit consists of three main components (Figure 1): (i) the core data capturing electronics based around a Raspberry Pi A+/B computer; (ii) a Huawei E3531 USB 3G mobile network dongle to enable continuous remote uploading of the data; and (iii) a solar powered battery system to provide a low-weight, renewable power source. Our open source software is available at www.github.com/sarabsethi/rpi-eco-monitoring and clear step-by-step setup and installation instructions are provided at www.rpi-eco-monitoring.com.

**Figure 1:**
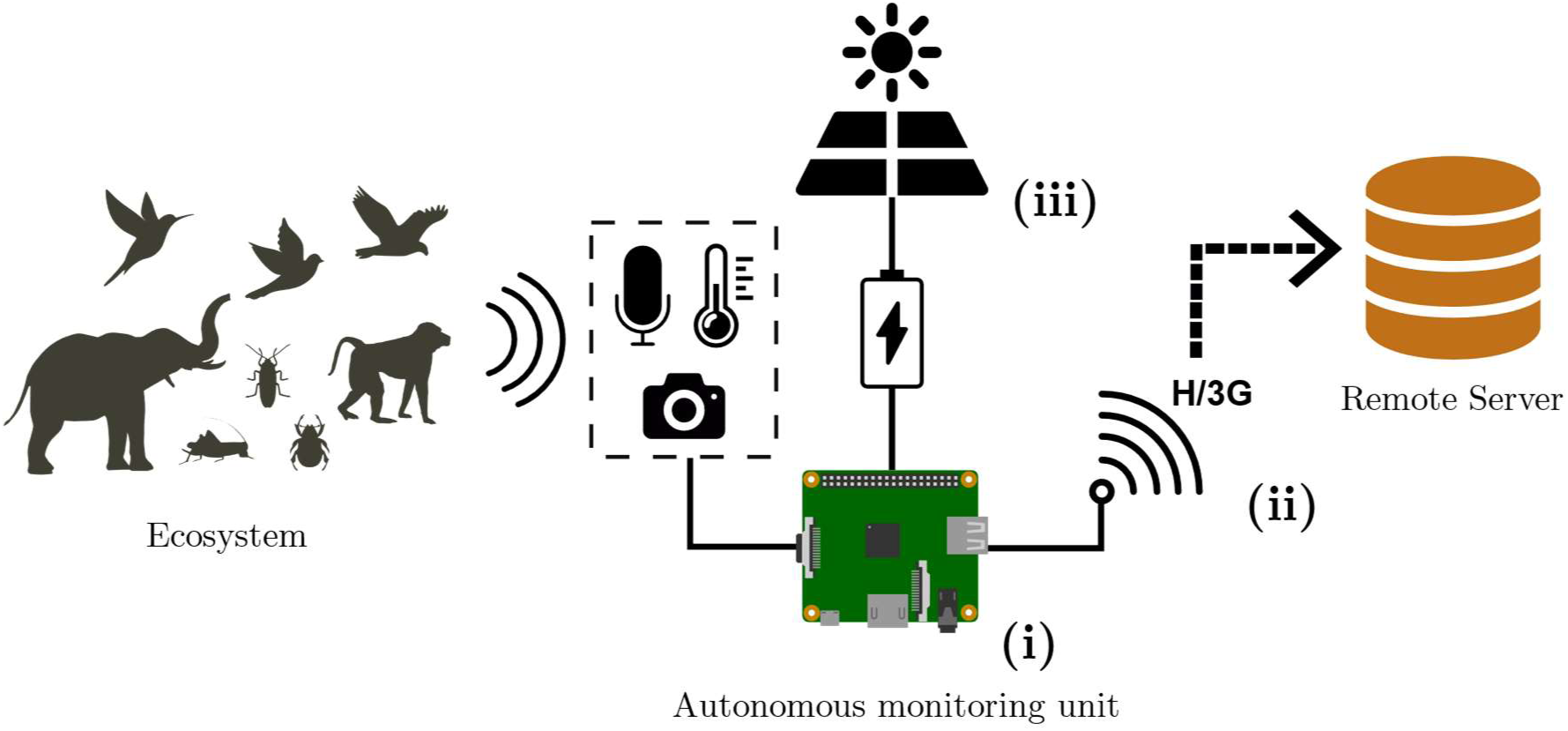
A schematic of the autonomous ecosystem monitoring system. Here we show how data is continuously captured from an ecosystem and uploaded automatically from the field to a remote server. The monitoring unit itself consists of three core components: **(i)** the core data capturing electronics, **(ii)** a mobile network link for uploading data, and **(iii)** a solar powered battery system.

The unit software is open-source Python and implements daily updates and logging to support remote management and trouble-shooting. The sensor software class separates data capture from post-processing and upload, and uses threading to allow these processes to run in parallel. We provide a clear generic class description, documenting the methods and setup needed to implement additional sensor types. Post-processed data files are stored in a local upload directory and transferred to a central FTP server via 3G. Local files are deleted after verified upload, hence the storage space on the device only acts as a buffer between the monitoring system and the remote server. This supports deployment in remote regions, where the mobile network signal can drop out for days at a time due to poorly maintained infrastructure. Our default set-up uses a 64GB micro-SD card to provide a high capacity buffer (e.g. ~1 month of compressed audio data, or 200 hours of uncompressed audio), but smaller, cheaper cards could be used in well-connected sites or for smaller data streams.

To ensure fully continuous monitoring over a long time period we use a solar powered battery system (Figure. 2). A Gamma 3.0 solar charge controller is connected to a 30W solar panel, a 12V 10Ah AGM (Absorbent Glass Mat) deep-cycle battery, and a 12V to 5V DC step down converter which ultimately powers the Raspberry Pi. When used for acoustic monitoring, the system draws approximately 1.5W, however this figure will vary significantly when using alternate sensors. The charge controller regulates the voltage coming from the solar panel and battery terminals to ensure a steady 12V DC is supplied to the load. In the case of low voltage, the sensor unit is shut down automatically to protect the battery from over-discharge and restarted when possible.

**Figure 2:**
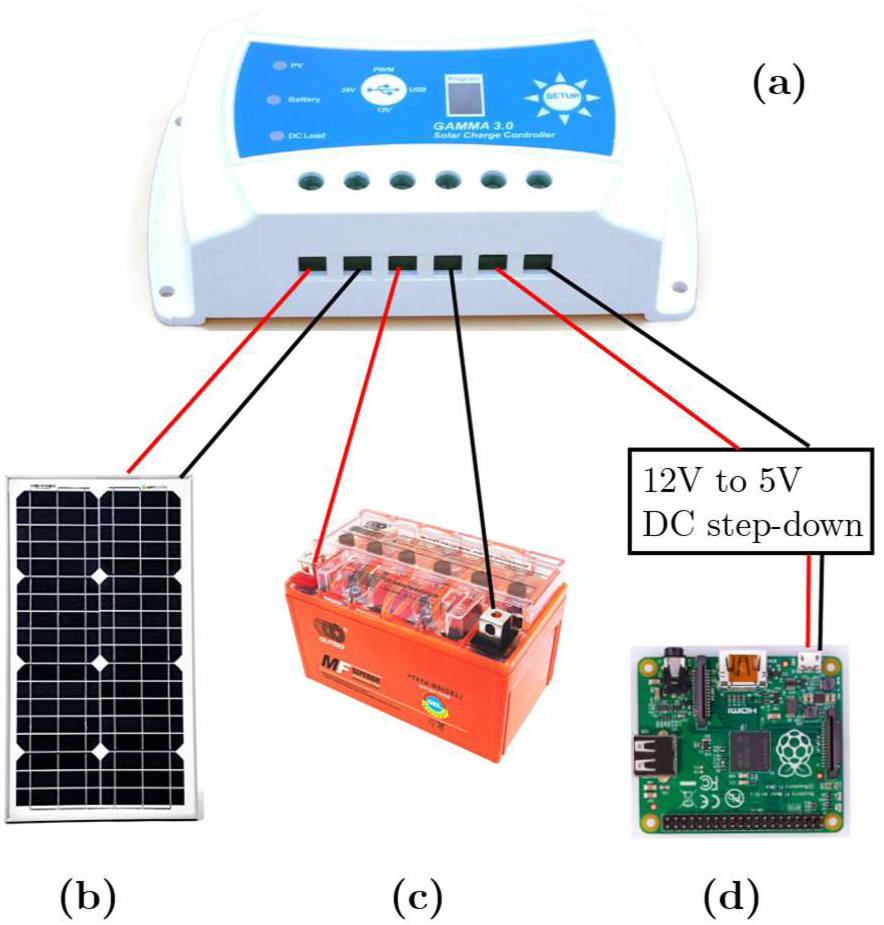
A schematic of the solar power system used. Here we show the solar power system used to power our recording unit continuously over long time periods. The components shown are: **(a)** a solar charge controller, **(b)** a 30W solar panel, **(c)** a 12V 10Ah AGM deep-cycle battery, and **(d)** the DC step down unit connecting the 12V load terminal to the 5V Raspberry Pi power input. Red and black lines indicate wires carrying positive and negative voltages respectively.

The total unit cost without sensors is around £180 (Supp. Table 1) and a 3G data plan is required for each unit (~ £30 per month, £360 per annum). With the addition of a £51 acoustic sensor package, this is considerably cheaper than closed-source commercial acoustic meters (e.g. Wildlife Acoustic Song Meter 4, ~£750) that do not typically have solar power or remote data upload and at least an order of magnitude cheaper than broadly comparable networked sensors (ARBIMON, ~£3000 excluding data link, Aide et al., 2013). The parts are generic and so can be easily optimised for other configurations, such as low data throughput sensors (smaller SD card), high power devices such as pH sensors (larger solar panel and battery) or more processor intensive data such as HD video (Raspberry Pi 3 B+).

## 3 Proof of concept

We have deployed these units in two configurations (continuous acoustic recording, Figure S1, and time lapse camera) in both tropical (Borneo) and temperate (UK) conditions to show viability in demanding physical conditions and in lower light environments. Our tropical field tests were based at the SAFE project, a large-scale fragmentation experiment located in the tropical rainforests of Sabah, Malaysia (Ewers et al., 2011). Deployment in rainforest requires that the packaging is robust to high temperature and humidity, and prevents insect ingress. Furthermore, forest canopy access is needed to ensure the solar panel is well illuminated.

Four initial deployments (starting from March 2017) demonstrated the importance of battery quality; sealed lead-acid (SLA) batteries are cheaper and more readily available but suffer from deterioration under high temperatures and frequent discharge cycling. Despite the battery deterioration, the monitoring units continued to boot and resume data recording automatically when enough power was available. Moreover, the remote uploading scheme also performed as designed, as all the files that were recorded on the monitoring units were successfully uploaded to the remote server despite regular network outages.

Since February 2017, we have deployed a further 11 units configured for continuous audio recording. In total, over 10,000 total hours of audio (approx. 630GB) has been recorded and transmitted from the SAFE project sites from these units. The longest fully continuous period of acoustic monitoring from the tropical forest recorders stands at 1,152 hours (48 days), after which the device stopped working during the nights (no daylight) and restarted the following morning. Due to supply issues all 11 recorders were using poor quality SLA batteries, and therefore suffered from regular outages as before. However, once again, the recording and remote uploading scheme resumed automatically and performed as designed when power was available. An acoustic monitoring unit set-up for testing in London in March 2016 has recorded and uploaded over 4,000 hours (approx. 250GB) of audio data. Additionally, we deployed two devices configured to capture time-lapse imagery, and we have received over 10,200 images (approx. 40GB) spanning a combined deployment time of 12 months.

Prolonged exposure to high temperature and humidity could have an adverse effect upon the electronics within our system. Through our first round of field tests we demonstrated that even after a six-month deployment in a tropical rainforest the autonomous monitoring system was able to record and remotely transmit data, having been exposed to temperatures ranging from approximately 2 to 31.5 degrees Celsius and an average 614mm of precipitation per month whilst within our rainforest sites.

## 4 Discussion

In this study we have introduced an inexpensive method of continuous, autonomous ecosystem monitoring. The equipment has been shown to be robust and versatile through successful field-testing in both tropical and temperate climates.

The volume of data collected through long-term continuous ecosystem monitoring using such equipment requires consideration. Other studies have performed analyses directly on the monitoring device at the point of data capture as a way of reducing the volume of data being transmitted (Deniz et al., 2017). This reduces the data storage requirements of the device since only analysed statistics derived from the raw data are kept. However, discarding the raw data limits the use of the monitoring unit as a general tool as it does not allow re-analysis of the original data using newer techniques. Furthermore, any analysis performed remotely on the recording device is likely to cause an increase in the power consumption.

Transmitting raw data does lead to important data storage considerations. 24 hour acoustic monitoring using VBR (Variable Bit Rate) MP3 compression leads to almost 700GB of data being recorded per unit in one year. Whilst many commercial providers offer large data storage options, plans on this scale can prove costly. Furthermore, performing real-time analyses on such a large volume of data will either place constraints on the complexity of the techniques used, or will require significant computing resources to be employed.

The requirement for a mobile data network obviously makes our solution unsuitable for the most remote locations, but in such situations the unit falls back to an offline mode and operates as a semi-autonomous recorder. Satellite internet and long-distance radio frequency links are proven technologies for long distance data collection but are expensive and require power. Adding mesh network capabilities to an array of our units might provide an alternative approach to gathering remote data to a single hub for storage or upload (Akyildiz and Wang, 2005; Dugas, 2005).

The efficiency of solar panels suffers significantly if the panels are not exposed to direct sunlight. We avoided this issue by installing the monitoring units in the canopy: the remote management of our units means that canopy access is only needed for deployment, retrieval and in case of malfunction. However, our design can easily be modified to use alternative power sources if required.

## 5 Conclusion

In this study we have outlined the design and implementation of an open-source, robust, autonomous ecosystem monitoring unit, which allows remote collection and transmission of field data over long time periods. The equipment can be built for significantly smaller costs if compared with existing systems offering comparative functionality, allowing field studies to be designed at a larger scale than previously possible. Because the hardware is open-source and low-cost, units can easily be replaced or ‘cannibalized’ for spares when inevitable damage occurs in the field. In addition to sharing the hardware design, we have open-sourced the code providing the reliable recording and upload mechanism, which should allow the broader scientific community to further develop this method for fully automated data collection.

## Acknowledgements

We would like to thank the research assistants and staff at the SAFE project camp, in particular Ryan Gray, Unding Jami, and the canopy access team. We would like to thank Henry Bernard for his collaboration. We acknowledge the support of the Imperial College Advanced Hackspace. This project was supported by funding from the Sime Darby Foundation and the Natural Environment Research Council [NE/K007270/1]. Sarab Sethi is also supported by Natural Environment Research Council through the Science and Solutions for a Changing Planet DTP.

## Authors’ Contributions

SS, RE, NJ, DO and LP contributed to the conceptualisation and final implementation of the study. SS and LP led the hardware design stage. SS and DO designed and wrote the open-source software. SS and RE conducted the field testing of the device. SS led the manuscript writing process with revisions provided by RE, NJ, DO and LP.

## Data Accessibility

All open source software and hardware design detailed in this study can be found at http://doi.org/10.5281/zenodo.1405359 (Sethi and Orme, 2018).

